# Mucus physically restricts influenza A viral particle access to the epithelium

**DOI:** 10.1101/2023.08.14.553271

**Authors:** Logan Kaler, Elizabeth M. Engle, Maria Rife, Ethan Iverson, Allison Boboltz, Maxinne A. Ignacio, Margaret A. Scull, Gregg A. Duncan

**Author notes:** Correspondence to: Gregg Duncan. These authors contributed equally to this work (Logan Kaler, Elizabeth M. Engle).

## Abstract

Prior work suggests influenza A virus (IAV) crosses the airway mucus barrier in a sialic acid-dependent manner through the actions of the viral envelope glycoproteins, hemagglutinin and neuraminidase. However, host and viral factors that influence how efficiently mucus traps IAV remain poorly defined. In this work, we assessed how the physicochemical properties of mucus influence its ability to effectively capture IAV using fluorescence video microscopy and multiple particle tracking. We found an airway mucus gel layer must be produced with virus-sized pores to physically constrain IAV. While sialic acid binding by IAV may improve mucus trapping efficiency, sialic acid binding preference was found to have little impact on IAV mobility and the fraction of viral particles expected to penetrate the mucus barrier. Further, we demonstrate synthetic polymeric hydrogels engineered with mucus-like architecture are similarly protective against IAV infection despite their lack of sialic acid decoy receptors. Together, this work provides new insights on mucus barrier function toward IAV with important implications on innate host defense and interspecies transmission.

## INTRODUCTION

Airway mucus is the first line of defense against inhaled particulates and pathogens.^1,2^ The mucus layer is comprised of mucins, which are heavily glycosylated gel-forming proteins.^3^ Mucin glycoproteins possess *O*-linked glycans that extend from the peptide backbone and are terminated with fucose, sialic acid, and sulfate.^4,5^ These mucin-associated glycans make up approximately 70–80% of the total mass of mucin and thus, play a critical role in the physical properties of mucus and its biological function in innate lung defense.^4–6^ Disulfide bonds between the cysteine-rich domains of the mucins and electrostatic interactions between mucin glycoproteins are responsible for the formation of the mucus gel network.^3^ The airway mucus gel effectively traps nano-scale particles and removes them via a process called mucociliary clearance, where cilia on the surface of the cell beat in coordination to move the mucus layer through the airway^3^. Prior work has demonstrated the functional benefits of the mucus barrier in preventing IAV infection. For example, infection by IAV was significantly inhibited in mice with lung-specific overexpression of the gel-forming mucin 5ac (Muc5ac) demonstrating its protective function.^7^ By mimicking the seasonal changes in humidity, prior work has also shown decreased air humidity dehydrates airway mucus leading to impaired mucociliary clearance and increased susceptibility to IAV infection in mice.^8^ Taken together, the mucus gel acts to directly block IAV and other viruses from entering the underlying epithelium by facilitating their clearance from the airway.

Prior studies on the mechanisms by which IAV overcomes the mucus barrier have primarily focused on the protective role of mucin-associated sialic acid (SA).^9^ IAV is an enveloped virus with two glycoproteins on the envelope that recognize SA: hemagglutinin (HA) and neuraminidase (NA).^2,9^ HA and NA work cooperatively to initiate infection in the airway epithelium, with HA binding SA while NA is responsible for solubilizing SA, favoring HA-SA dissolution.^9^ Glycans containing α-2,3 and α-2,6 linked SA are found on the surface of the epithelial cells in the respiratory tract as well as on mucins. In prior work, it was found that inhibition of NA leads to immobilization of IAV particles in airway mucus.^10,11^ In addition, chemical and/or enzymatic removal of mucin-associated SA has been shown to enhance the mobility of IAV through mucus.^10,11^ This suggests mucin sialylation enables mucus gel trapping of IAV through direct binding by HA and release of SA by NA enables IAV to efficiently bypass the mucus barrier.

However, this past work did not fully consider the physical constraints imposed by mucus as a hydrogel with pores ranging in sizes from 100-500 nm on IAV particles with dimensions ranging from ∼120 nm in a spherical form to ≥250 nm in a filamentous form. ^3,12–14^ In previous work from our group, we observed IAV diffused in human mucus at a similar rate to synthetic, muco-inert nanoparticles with a diameter comparable to IAV.^15^ This would suggest the mucus barrier acts to physically block the penetration of IAV rather than adhesively trap viral particles through IAV binding to SA. It should be noted that another study also observed little evidence of SA-mediated trapping of IAV within mucus and alternatively proposed that neutralizing antibodies against IAV facilitate entrapment in the mucus barrier.^16^ Considering these past observations by our group and others, the features of mucus that render it permissive to IAV particles has yet to be clearly established. Further, the role of IAV binding preference for α2,3-or α2,6-SA on virus trapping within mucus is largely unaddressed. We also note prior reports have shown airway mucins in a soluble form can competitively inhibit infection by IAV and other viruses.^17,18^ We consider these direct antiviral effects as distinct from mucus gel barrier functions that facilitate capture and removal of viral particles from the airway.

In this work, we used A/Udorn/307/72 (Udorn), a H3N2 IAV that possesses pleomorphic particle morphologies with both spherical and filamentous shape, to study how IAV navigates through airway mucus with physically and biochemically distinct barrier properties. To study the impact of sialic acid preference on IAV penetration through mucus, we evaluated the diffusion of Udorn IAV with mutations to specific residues in the receptor binding domain that alter its preference to either α2,3-or α2,6-SA.^19,20^ Our previous study was conducted using *ex vivo* human airway mucus collected from patients to evaluate IAV diffusion through mucus.^15^ However, the properties of mucus samples vary significantly from patient-to-patient. To provide a source of mucus with consistent properties, we harvested mucus from tissue cultures consisting of 2 human airway epithelial cell (HAE) lines, in addition to normal human bronchial epithelial (NHBE) primary cells for this work. Mucus collections from these three different lung epithelial cell sources allowed us to compare their ability to trap IAV and how this may relate to their biomolecular properties. To interrogate the impact of the physical barrier function of mucus on infection, we compared the extent of infection of Udorn IAV in HAE cultures coated with a protective mucus layer or an engineered hydrogel layer with similar pore network size but lacking any glycans to serve as adhesive sites for IAV particle trapping.

## RESULTS

### Biochemical characterization of human lung epithelial cell-derived mucus

To study the mobility of IAV within human airway mucus, lung epithelial cells grown at an air-liquid interface (ALI) were used to generate mucus which could be regularly collected for experimental use. Mucus collected from Calu-3, BCi-NS1.1 (BCi),^21^ and primary NHBE cultures was characterized for relative mucin content, disulfide bond (cystine) concentration, and SA content using fluorometric assays **(Fig. 1A-C)**. NHBE mucus possessed significantly higher concentrations of relative mucin in comparison to both BCi and Calu-3 mucus (**Fig. 1A**). In comparison to Calu-3 mucus, relative mucin content was ∼1.8-fold higher in BCi mucus and ∼4.3-fold higher in NHBE mucus pooled across four donors. Conversely, Calu-3 mucus was found to possess significantly higher disulfide bonds per unit mass mucin in comparison to BCi and NHBE mucus (**Fig. 1B)**. There were no significant differences in the disulfide bond concentration per unit mass mucin between BCi and NHBE mucus. We also found the concentration of SA per unit mass mucin in NHBE mucus (72 ± 11 mM / g mucin) was ∼2.7-fold higher than that found in BCi mucus (27 ± 0.3 mM / g mucin) and ∼1.4 fold higher than that found in Calu-3 mucus (51 ± 9 mM / g mucin) (**Fig. 1C)**.

**Figure 1.**
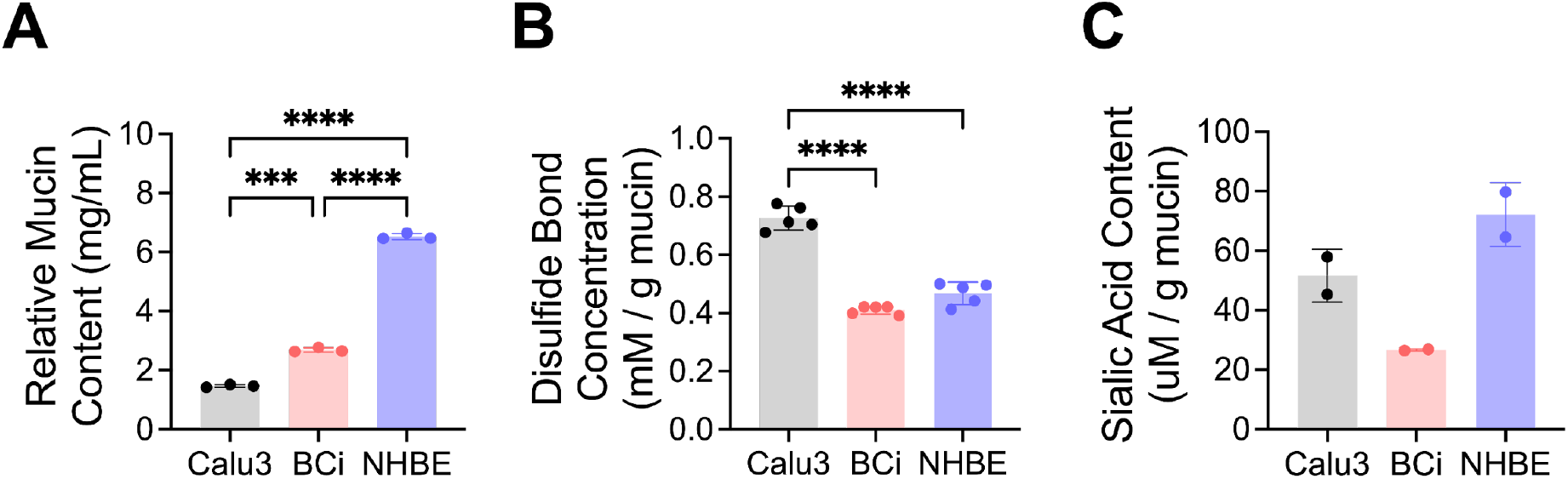
Biochemical characterization of human lung epithelial cell-derived mucus. **(A-C)**Characterization of Calu-3, BCi, and NHBE mucus to determine **(A)** relative mucin content, **(B)** disulfide bond concentration per unit mass mucin, and **(C)** SA content normalized per unit mass mucin. Bars indicate mean with error bars for standard deviation with measurements from each technical replicate shown as points. Data set in (**A**,**B**) statistically analyzed with one-way ANOVA and Šídák’s multiple comparisons test: ns = not significant; **p < 0.01, ****p < 0.0001.

### Udorn IAV and virus-sized nanoparticle diffusion in airway mucus

The diffusion of muco-inert nanoparticles (NP) and fluorescently-labeled Udorn IAV was assessed in mucus harvested from Calu-3, BCi, and NHBE ALI cultures **(Fig. 2A)**. We confirmed muco-inert NP and Udorn IAV particle sizes were in a similar range based on dynamic light scattering measurements (**Fig. S1**). Thus, muco-inert NP serve as an experimental control that indicate the potential impact of mucus microstructure on IAV trapping. Diffusion rates measured for both NP and Udorn IAV were determined in the same regions of interest using fluorescence video microscopy. Multiple particle tracking analysis of NP indicated the resulting diffusion rate of NP, as measured by the mean squared displacement at a time scale of 1 second (MSD_1s_), was significantly higher in Calu-3 mucus in comparison to BCi and NHBE mucus **(Fig. 2B)**. Based on measured NP diffusion, we estimated the pore size (ξ) of the mucus network **(Fig. 2C)**. For comparison to the estimated mucus pore sizes, the size range of Udorn IAV particles measured by dynamic light scattering is also highlighted in gray. Calu-3 mucus possessed the largest pores ranging from 1000–2000 nm (**Fig. 2C**). In comparison, BCi and NHBE mucus pore sizes were much smaller with values ranging from approximately 200–900 nm and 350–950 nm, respectively. Particle tracking analysis showed Udorn particle diffusion was significantly increased in Calu-3 mucus compared to both NHBE and BCi mucus **(Fig. 2D)**.

**Figure 2.**
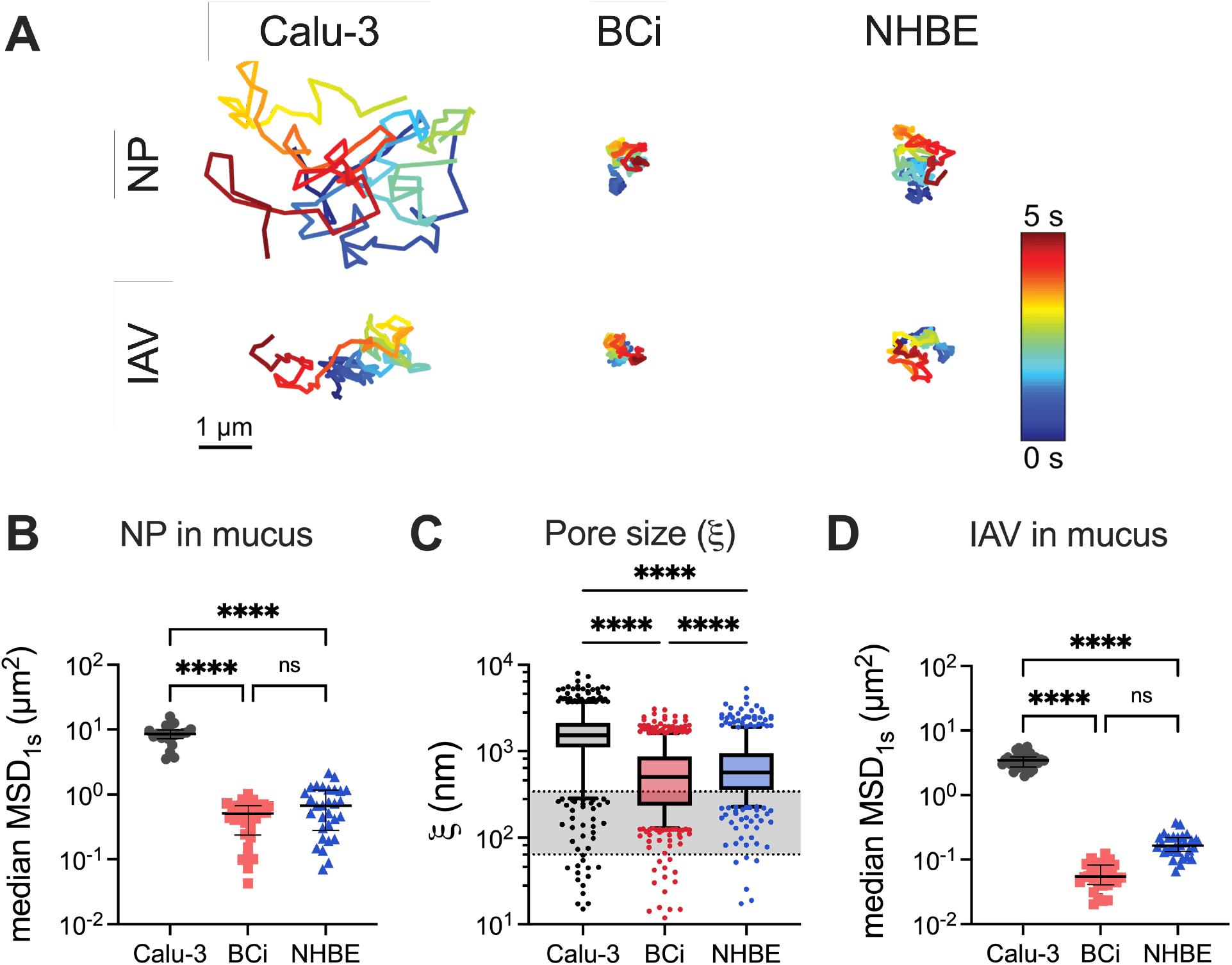
Udorn IAV and virus-sized nanoparticle diffusion in airway mucus. **(A)** Representative trajectories for diffusion of NP and Udorn in Calu-3, BCi, and NHBE mucus. Trajectory color changes with time, with dark blue indicating 0 s and dark red indicating 5 s. Scale bar = 1 μm. **(B)** Measured MSD at time scale of 1 second (MSD_1s_) for NP diffusion in Calu-3, BCi, and NHBE mucus. Each data point represents the median measured MSD_1s_ in each video with at least 5 videos from 4-6 technical replicates. Black lines indicate overall median MSD_1s_ and brackets indicate interquartile range. **(C)** Estimated pore size (ξ) from NP diffusion in Calu-3, BCi-NS1.1, and NHBE mucus. Shaded region indicates size range of Udorn particles as measured by dynamic light scattering. **(D)** Median MSD_1s_ for Udorn IAV diffusion in Calu-3, BCi, and NHBE mucus. Data sets in (**B**,**D**) analyzed with one-way analysis of variance (ANOVA) and Šídák’s multiple comparisons test: ns = not significant; ****p < 0.0001. Data sets in (**C**) statistically analyzed using Kruskal-Wallis test with Dunn’s test for multiple comparisons: ****p < 0.0001.

### Impact of sialic acid depletion on IAV diffusion through mucus

To account for the role of SA cleavage on IAV diffusion, we evaluated the extent to which mucin-associated SA was removed by Udorn. NHBE mucus was used for these studies given the higher abundance of SA found in these samples compared to other mucus sources (**Fig. 1B**). Following 30 minutes of incubation, we found IAV treated mucus had significantly reduced total SA concentration (**Fig. 3A**). Similarly, introduction of an exogenous NA (NA_ex_) from *Arthrobacter ureafaciens* was introduced to hydrolyze terminal SA.^22^ Treatment with NA_ex_ resulted in an 8-fold decrease in SA concentration compared to untreated mucus **(Fig. 3B)**. Multiple particle tracking was then used to analyze NP and Udorn movement in untreated and NA_ex_ treated mucus (**Fig. 3C**). Measured MSD_1s_ showed that Udorn IAV particles moved at a similar rate in NA_ex_ treated mucus compared to untreated mucus. We also found NP diffused significantly faster than Udorn in untreated NHBE mucus. However, there was no significant difference in IAV and NP diffusion in NA_ex_ treated mucus.

**Figure 3.**
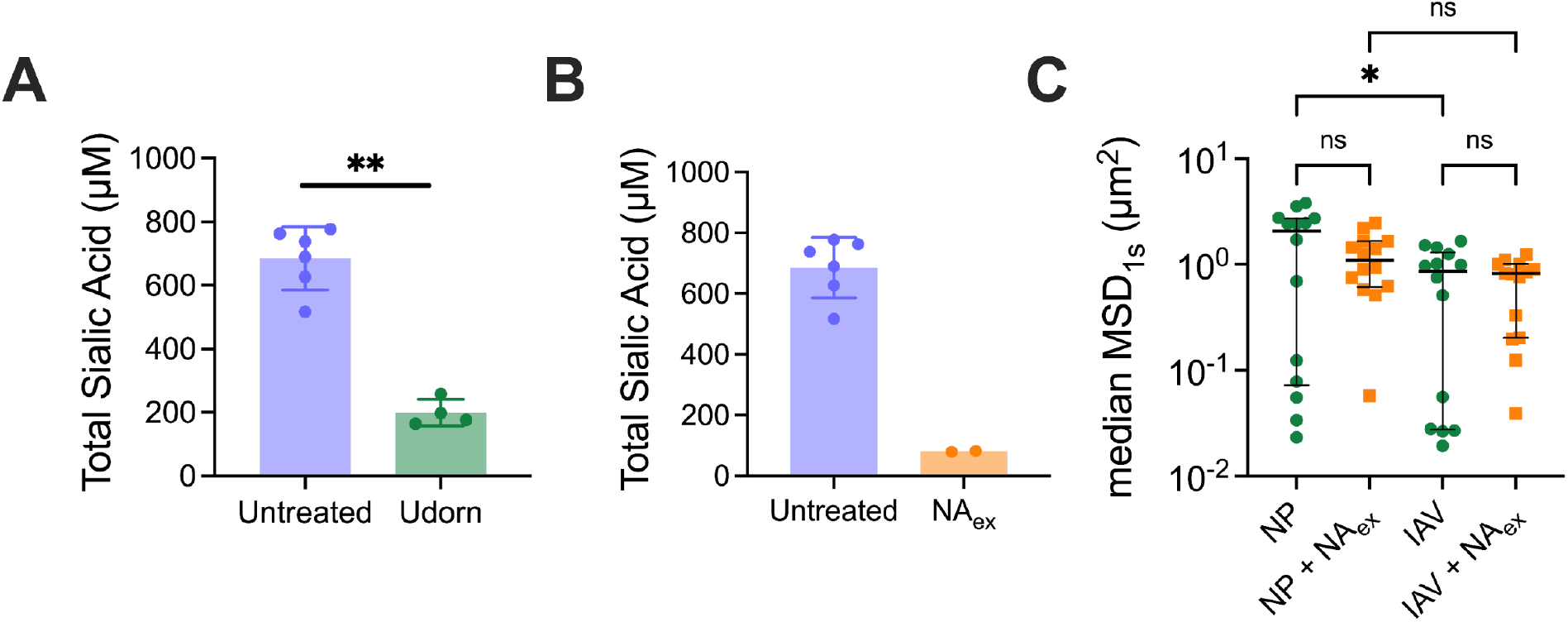
Impact of sialic acid depletion on IAV diffusion through mucus. Characterization of total SA concentration for (**A**) Udorn IAV treated and (**B**) NA_ex_ treated NHBE mucus. Data set in **(A)** statistically analyzed with an unpaired T-test: **p < 0.01. Trajectory color changes with time, with dark blue indicating 0 s and dark red indicating 10 s. Scale bar = 1 μm. **(C)** Median MSD_1s_ for NP and Udorn in untreated and NA_ex_ treated mucus. Black line indicates overall median MSD_1s_ and brackets indicate interquartile range. Data set in **(C)** analyzed with one-way analysis of variance (ANOVA) and Šídák’s multiple comparisons test: ns = not significant; *p < 0.05.

### Impact of SA preference on IAV penetration through the mucus barrier

To probe the impact of SA binding preference on IAV diffusion, we used two previously established Udorn mutants that possess an HA with preferential binding for either α2,3-SA (Ud23) or α2,6-SA (Ud26).^20^ Diffusion of each Udorn mutant in NHBE mucus was determined in the same regions of interest as wildtype Udorn and muco-inert NP. Based on particle tracking analysis of wildtype and mutant Udorn in NHBE mucus (**Fig. 4A)**, measured MSD_1s_ indicates a similar diffusivity for Udorn when compared to Ud23 and Ud26 **(Fig. 4B**,**C)**. We also found NP were significantly more diffusive than Udorn, Ud23, and Ud26 IAV particle types. Wildtype and mutant Udorn IAV diffusion were also evaluated in mucus harvested from Calu-3 and NHBE where similar trends were observed with minimal changes in diffusivity between virus types (**Fig. S2**).

**Figure 4.**
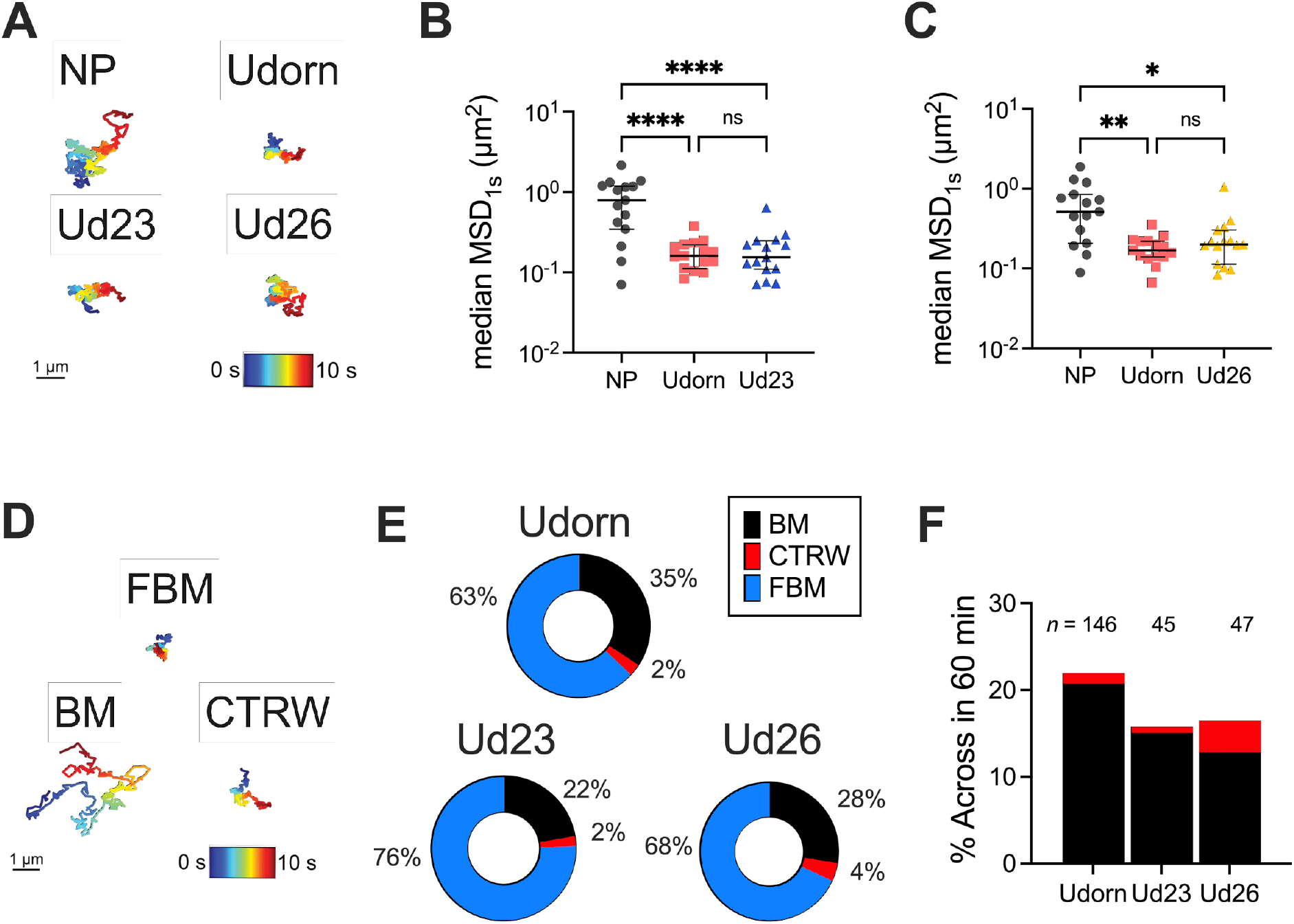
Impact of SA preference on IAV penetration through the mucus barrier. **(A)** Representative trajectories for NP, Udorn, Ud23, and Ud26 diffusing in NHBE mucus. Trajectory color changes with time, with dark blue indicating 0 s and dark red indicating 10 s. Scale bar = 1 μm. **(B)** Median MSD_1s_ for NP, Udorn, and Ud23 diffusing in NHBE mucus. **(C)** Median MSD_1s_ for NP, Udorn, and Ud26 diffusing in NHBE mucus. In **(B-C)**, NP is shown as gray circles, Udorn is shown as red squares, Ud23 is shown as blue triangles, and Ud26 is shown as yellow triangles. Black line indicates overall median MSD_1s_ and brackets indicate interquartile range. Data sets in **(B**,**C)** statistically analyzed with one-way ANOVA and Šídák’s multiple comparisons test: *p < 0.05, **p < 0.01, ****p < 0.0001. **(D)** Representative trajectories of Udorn exhibiting FBM, BM, and CTRW diffusion in NHBE mucus. **(E)** Normalized percent of wildtype and mutant Udorn exhibiting FBM, BM, and CTRW motion. **(F)** Percent of wildtype and mutant Udorn particles to cross the 10 μm mucus barrier in 60 minutes with the corresponding total number of particles for each sample. In **(E-F)**, BM is shown as black bars, and CTRW is shown as red bars.

To gain further insight on the dependence of IAV transport through mucus on SA binding preference, we computationally predicted the time required for each IAV type to penetrate a 10 μm-thick mucus barrier using a machine learning-based approach developed in prior work.^23^ First, machine learning was used to classify individual IAV trajectories as either subdiffusive or diffusive. The subdiffusive particles were further subdivided as exhibiting either fractional Brownian (FBM) or continuous time random walk (CTRW) motion.^24^ FBM particles follow a random walk, but the following step has a higher probability to be in the opposite direction than the previous step.^24^ CTRW particles are characterized by random jumps in time and space, resulting in a “hopping” motion.^24^ Diffusive particles are undergoing Brownian motion (BM), which is classical thermally driven diffusion, characterized as a random walk with steps taken to the left and right with equal probability.^25^ The resulting classification of particles indicated the majority of Udorn, Ud23, and Ud26 particles were exhibiting FBM with a smaller percentage of BM **(Fig. 4D-E)**. Interestingly, all IAV strains had some percentage of particles that exhibited CTRW “hopping” movement in NHBE mucus, with Ud26 having the largest percentage of CTRW particles. Based on individual diffusion modes, the percentage of particles to cross the mucus barrier in 60 minutes was mathematically predicted for each Udorn IAV type **(Fig. 4F)**. Udorn was predicted to have the largest percentage of particles across the mucus barrier in 60 minutes compared to all other IAV types in NHBE mucus, with ∼22%, while Ud23 and Ud26 have predicted percentages of ∼16% **(Fig. 4F)**. For all IAV types, particles exhibiting FBM were not predicted to cross the mucus barrier in a physiological timeframe.

### IAV infection in human airway epithelial cultures with mucus and PEG gel coatings

To further assess the contributions of pore network structure within mucus to its ability to effectively capture viral particles and prevent infection, a synthetic hydrogel was developed as a model system for direct comparison to mucus as a protective barrier against IAV. Specifically, we engineered a polyethylene glycol (PEG)-based hydrogel which readily forms upon mixing via disulfide bond formation (**Fig. 5A**). We found PEG gels composed of 2% w/v thiolated 10 kDa 4-arm PEG and 2% w/v orthopyridyl disulfide (OPSS) functionalized 5 kDa 4-arm PEG formed into a gel with pore sizes with similar dimensions to naturally-secreted BCi mucus (**Fig. 5B**). We next evaluated the IAV trapping capacity where we found a significant reduction in the diffusion rate of Udorn IAV within the PEG gel as compared to BCi mucus (**Fig. 5C**). With this established, we continued with this model hydrogel to determine if its capacity for physical capture of IAV would render it an effective barrier against IAV infection. A diagram outlining the workflow for the IAV challenge experiments is provided in **Fig. 5D**. To allow for comparison of the protective function of BCi mucus and PEG gels, mucus was harvested from differentiated BCi HAE cultures. Prior to challenge with IAV, BCi HAE cultures are washed to remove endogenous mucus and BCi mucus or PEG gel precursor solution was apically applied to evenly coat the epithelial surface We confirmed PEG gel treatment did not have an impact on cell viability and produced a gel thickness within a physiological range of ∼50 μm (**Fig. S3**). Once equilibrated, cultures were inoculated with a low concentration of IAV and washed to remove inoculum after 15 minutes. All cultures were subsequently treated with zanamivir (1.25 μM) to prevent secondary infections. Based upon staining for IAV nucleoprotein, we observed a significant reduction in infection by IAV in cultures coated with either BCi mucus or PEG gel compared to washed (uncoated) controls (**Fig. 5E-H**). These studies were also repeated in the absence of zanamivir and reduced infection was also observed in HAE cultures coated with either BCi mucus or PEG gel (**Fig. S4**).

**Figure 5.**
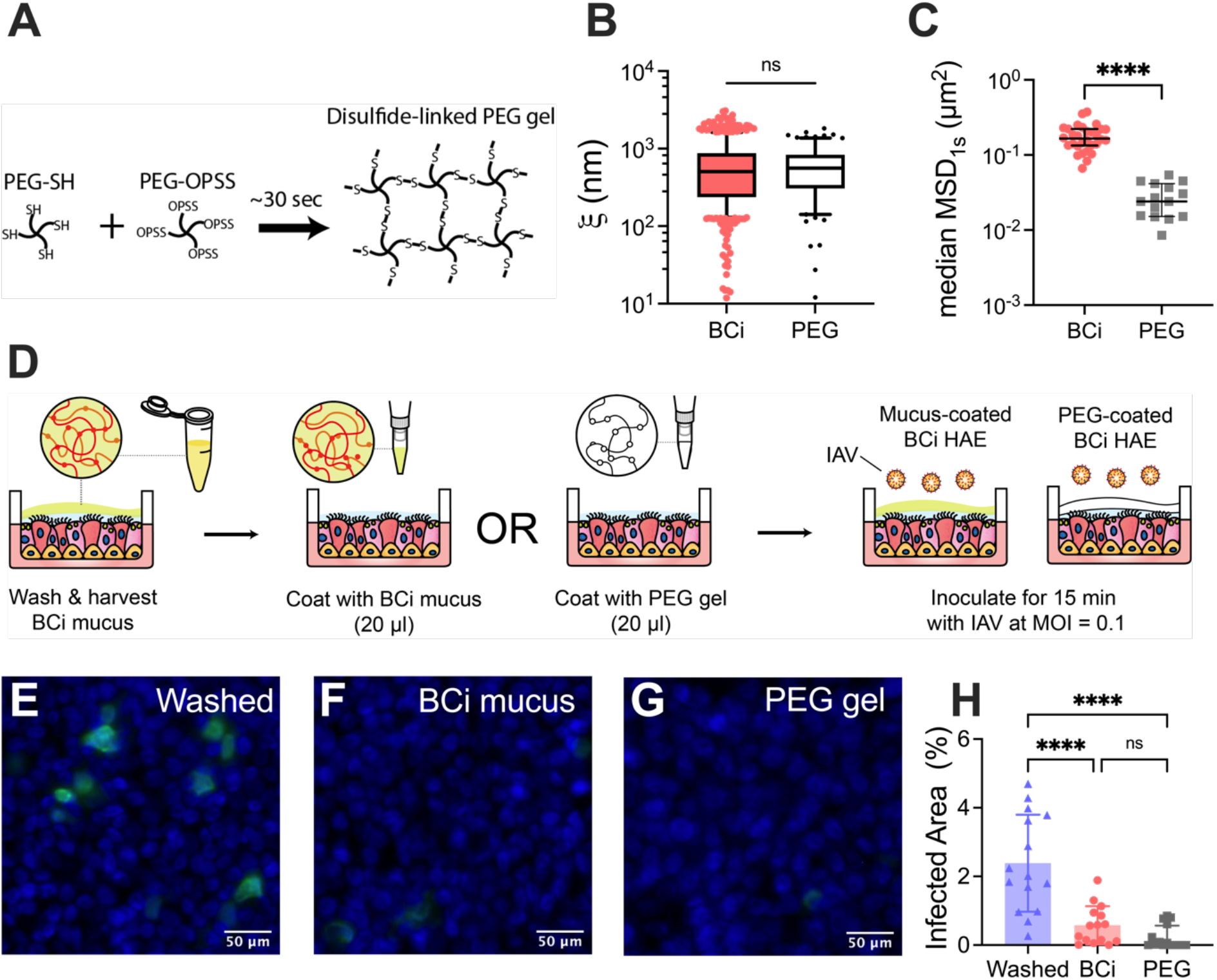
IAV infection in human airway epithelial cultures with mucus and synthetic PEG gel coatings. (**A**) PEG hydrogels were prepared using 10 kDa thiol-modified and 5 kDa OPSS-modified 4-arm PEG polymers which rapidly form disulfide bonds upon mixing. (**B**) Estimated pore size (ξ) based on NP diffusion in BCi mucus and PEG gels. Data set in **(B)** statistically analyzed with Kruskal-Wallis test with Dunn’s test for multiple comparisons: ns = not significant. (**C**) Measured log_10_[MSD_1s_] for Udorn in BCi mucus and PEG gels. Data set in **(C)** statistically analyzed with one-way ANOVA and Šídák’s multiple comparisons test: ****p < 0.0001. (**D**) Schematic illustrating protocol to introduce BCi mucus and PEG gels as barriers to BCi human airway epithelial (HAE) cultures to determine their impact on infection. (**E-G)** Fluorescence micrographs of (**E**) washed, **(F)** BCi mucus coated, and (**G)** PEG gel coated HAE cultures infected apically with Udorn IAV 12 hpi after 15 min of inoculation. Green indicates staining for IAV nucleoprotein and blue indicates DAPI-stained nuclei. Scale bar = 50 μm. **(H)** Percentage of HAE culture area infected as determined by IAV nucleoprotein staining. Each data point representative of individual fluorescent micrographs collected from 3 individual cultures per experimental condition.

## DISCUSSION

In this work, we used video microscopy and multiple particle tracking analysis to evaluate how size-limited transport and adhesive SA binding influence mucus barrier function towards IAV. We conducted these studies using mucus harvested from 3 commonly used lung epithelial cell culture models. Our studies revealed significant differences in the biochemical and biophysical properties of mucus produced by each cell type. Based on our measurements of NP diffusion, Calu-3 mucus possesses pore sizes on the order of microns which is far larger than what has been previously reported for human airway mucus collected *ex vivo*.^14,26,27^ This is likely explained by the lower mucin content of Calu-3 mucus which falls below the overlap concentration of 2-4 mg/mL previously determined for mucins in a semi-dilute concentration regime.^28^ At or above this overlap concentration, the spacing between neighboring mucin polymers is minimized to facilitate intermolecular (noncovalent) interactions to stabilize the mucus gel.^28^ Based on measured mucin content (∼1.5 mg/mL), Calu-3 mucus would be in a dilute concentration regime leading to reduced mucin-mucin interactions and greater mucus gel porosity. We also observed a tighter network structure in BCi mucus as compared to NHBE mucus. Based on the measured mucin concentration, we would have predicted a denser network to form in NHBE mucus in comparison to BCi mucus. However, this observation may be explained by differences in mucin glycosylation and/or mucin subtypes (e.g. MUC5B, MUC5AC) present in each gel type. For example, we have previously observed mucus gels composed of MUC5AC possess a smaller pore size in comparison to mucus gels composed of MUC5B.^29^ Additional biochemical analyses will be needed in future work to explain these differences.

Our measurements of Udorn IAV diffusion in each mucus type revealed gels with smaller pores were more effective at virus trapping. This can be explained intuitively as mucus gels with narrower openings between mucin fibers can physically obstruct IAV diffusion. Based on the measured size range of Udorn IAV, we estimate approximately 7%, 40%, and 24% of the measured pore sizes for Calu-3, BCi, and NHBE mucus, respectively, are small enough in size to sterically hinder diffusion and physically capture Udorn IAV within the mucus gel. This explains the observed differences in IAV diffusion in different mucus sources where BCi and NHBE mucus, containing a larger fraction of virus-sized pores, posed the greatest hindrance to IAV mobility. We also found mucus derived from BCi and NHBE cultures possessed similar capacity to trap IAV despite the greater than 2-fold difference in mucin-associated SA content. To assess the direct impact of physical trapping of IAV particles on infection, we generated PEG hydrogels that emulated the physical structure of BCi mucus for use as an artificial mucosal barrier. While both the PEG and BCi mucus gels had similar pore sizes, we found IAV diffusion was significantly reduced within the PEG gel in comparison to BCi mucus. This may be explained by the more homogenous network structure of the PEG gel as compared to BCi mucus which is more complex in composition leading to greater variation in its internal structure. We also found application of the PEG gels and BCi mucus to HAE cultures provided similar protection against Udorn IAV infection. This indicates the mucus-like architecture of the PEG gels posed a significant barrier to infection even in the absence of sialic acid decoy receptors to mediate IAV trapping. Taken together, our data suggests mucus network size can significantly impact IAV mobility and successful initiation of infection.

Consistent with prior reports,^10,11,30^ we observed Udorn IAV was capable of partially depleting the mucus barrier of sialic acid which may limit their ability to act as decoy receptors and influence IAV mobility. Accordingly, we also observed the enzymatic depletion of SA with exogenous NA did not lead to enhanced IAV mobility. Given IAV’s natural ability to remove SA decoy receptors, removal of mucin-associated SA via enzymatic treatment may not facilitate significantly greater penetration of IAV particles through the mucus barrier. However, unlike untreated controls, IAV diffused at a similar rate to muco-inert NP following NA_ex_ treatment of NHBE mucus. This is likely explained by a reduction in IAV-mucus binding due to the loss of SA receptors that can help to slow IAV diffusion. We also used a previously reported approach to chemically remove Sia^11^ from NHBE mucus that involves sodium periodate (NaIO_4_) treatment to oxidize the bonds between adjacent hydroxyls of mucin-associated sugars (**Fig. S5**). This resulted in a 2.3-fold decrease in SA concentration, which was similar to the reductions in SA as mucus treated with Udorn alone. In addition, there was an apparent increase in NP diffusion following NaIO_4_ treatment indicative of an increase in mucus pore size. Thus, it is unclear from these data if the resulting increases in IAV diffusion can be attributed solely to the impacts of NaIO_4_ treatment on mucin-associated SA.

To evaluate the importance of SA binding preference, we utilized two Udorn mutants that preferentially bind either α2,3-or α2,6-SA. We hypothesized Ud23 viral particles would be more readily trapped due to a higher concentration of α2,3-SA linkages in airway mucus as determined in previous work.^31–33^ However, there were no significant differences in the diffusion rates of Udorn, Ud23, and Ud26 IAV in NHBE mucus. This may also be attributed to the intrinsic activity of Udorn NA and cleavage of α2,3-and/or α2,6-SA within the mucus gel. In addition, we predicted the highest total percentage of Udorn particles would cross the mucus barrier in the NHBE mucus. Interestingly, very similar fractions of Ud23 and Ud26 IAV particles are predicted to bypass the mucus barrier. These data suggest SA preference has a relatively small impact on the protective functions of the mucus barrier against IAV. Udorn, Ud23, and Ud26 IAV diffusion was also evaluated in Calu-3 and BCi mucus where we observed increased diffusivity in Calu-3 compared to BCi mucus. This can likely be attributed to the Calu-3 mucus network possessing the largest pore size, as indicated by the muco-inert NP.

As we consider the potential implications of this work, we acknowledge our study had several limitations. The relative concentrations of α2,3-vs α2,6-SA terminated mucin glycans were not quantitatively assessed in the mucus collected for these studies and this is likely to influence the results of our studies. The Udorn mutants used in this work possessed alterations in SA preference for HA envelope proteins. Inclusion of a pseudo-typed IAV with matched NA and HA pairs with preference for α2,3-vs α2,6-linked SA could give additional insights in future work. Further, inactivation of NA via mutation would allow for the analysis of the effect of HA activity on mobility independent of NA activity in future studies. We are also keenly aware the direct antiviral functions of airway mucins, which as noted previously are not considered here, may rely on SA-terminated glycans in the mucus gel which could facilitate competitive inhibition of IAV binding to the airway epithelium. Furthermore, partitioning of IAV particles at the air-mucus interface may also be facilitated by SA binding which this work does not address.

The findings of this study agree with our previous work indicating that IAV mobility is dependent on the network size of the mucus, as opposed to IAV-mucus binding alone.^15^ Overall, our data emphasizes the role of network size rather than glycan-specific interactions as responsible for IAV movement through the mucus barrier. We also discovered SA binding preferences had less of a role in IAV-mucus interactions which could be due to NA-mediated SA release. The extent of IAV binding to the mucus gel is also likely to be influenced by the presence of primarily *O*-linked SA in mucin, as opposed to primarily *N*-linked SA on the surface of airway epithelial cells.^34^ Direct profiling of HA binding to mucins and *O*-linked mucin glycan is likely needed to determine mucin-associated SA preferences for IAV. We also note the glycan profile of the mucus barrier may be altered between individuals, as a function of age, and as a result of underlying lung disease.^35,36^ Alterations to mucin glycans and their impacts on the mucus barrier towards IAV and other respiratory viruses should also be considered in future work. Overall, this work provides further insight into the functional role of mucus in IAV pathogenesis and zoonotic transmission.

## ACKNOWLEDGEMENTS

This project was funded by the Burroughs Wellcome Fund CASI (to G.A.D.), Cystic Fibrosis Foundation (BOBOLT23H0 to A.B.), the National Institutes of Health (R21 AI142050 to M.A.S. & G.A.D., R01 HL160540 to G.A.D., R01 HL151840 to M.A.S., T32 AI089621 to L.K., M.R., E.I.), and the National Science Foundation (CBET 2129624 to G.A.D & M.A.S.).

## AUTHOR CONTRIBUTIONS

L.K. and E.M.E. conceived, designed, and performed the research and led data analysis. E.I. produced viruses used for this study. A.B, M.A.I, and M.R. helped with experiments including human airway epithelial cell mucus collection and viral challenge studies. M.A.S. and G.A.D. conceived and designed experiments. L.K., E.M.E. and G.A.D. wrote the article. All authors reviewed and edited the article.

## DECLARATION OF INTERESTS

The authors declare no competing interests.

## MATERIALS AND METHODS

### Cell Culture

Human lung adenocarcinoma cells (Calu-3 cells) were purchased from ATCC (HTB-55). The immortalized HAE line BCi-NS1.1 was kindly provided by Matthew Walters and Ronald Crystal (Weill Cornell Medical College)^21^. Human airway tracheobronchial epithelial cells (NHBE) isolated from airway specimens from four donors without underlying lung disease were provided by Lonza, Inc. The donor information for the NHBE cells is included in **Supplemental Table 1**. BCi-NS1.1 and NHBE cells were first expanded on plastic in Pneumacult-Ex or Pneumacult-Ex Plus medium (no. 05008 or 05040, StemCell Technologies). Airway cells were then seeded (3.3 × 10^4^ cells/well) on rat tail collagen type 1-coated permeable Transwell membrane supports (6.5 mm; no. 3470, Corning, Inc.) and differentiated in Pneumacult-ALI medium (no. 05001, StemCell Technologies) with provision of an air-liquid interface (ALI) for at least 28 days until matured into a pseudostratified mucus-producing and ciliated epithelium. Calu-3 cells were expanded and maintained in Eagle’s Minimum Essential Medium (EMEM; ATCC) with 10% fetal bovine serum (FBS; Sigma-Aldrich) and 1% Penicillin-Streptomycin solution (Pen-Strep; Sigma-Aldrich). Cells were seeded on collagen-treated (Sigma Aldrich) PET 0.4 μm 24-well hanging inserts (EMD Millipore) for air-liquid interface (ALI) culturing. Cells were maintained at ALI for 25 days to allow for polarization and mucus production. All cell cultures were maintained at 37°C with 5% CO2.

### Virus Strains

The reverse genetics system for influenza A virus A/Udorn/307/72 (Udorn) was a gift from Robert Lamb. Infectious virus was rescued from cloned cDNAs in 293T and MDCK cells as previously described.^37^ Two Udorn mutants were prepared according to previous works: Udorn with an HA mutation for preferential α2,3-SA binding (Ud23), and Udorn with an HA mutation for preferential α2,6-SA binding (Ud26)^20,38^. Ud23 has two mutations in the receptor binding domain, L226Q and S228G, while Ud26 has a single mutation in the receptor binding domain, E190D20. Udorn IAV was labeled with a lipophilic dye, 1,1′-dioctadecyl-3,3,3′,3′-tetramethylindocarbocyanine perchlorate (DiI; Invitrogen) while Ud23 and Ud26 IAV were labeled with a different lipophilic dye with a longer excitation and emission wavelength, 1,1′-dioctadecyl-3,3,3′,3′-tetramethylindodicarbocyanine 4-chlorobenzenesulfonate salt (DiD; Invitrogen). IAV sizes were determined via dynamic light scattering (DLS) using the NanoBrook Omi (Brookhaven Instruments).

### Nanoparticle preparation

As previously described, carboxylate modified fluorescent polystyrene nanoparticles (NP; Life Technologies) with a diameter of 100 nm were coated with 5 kDa methoxy polyethylene glycol (PEG)-amine (Creative PEGWorks) via a carboxyl-amine linkage to generate muco-inert nanoparticles.^39^ NP size was determined via DLS using the NanoBrook Omi (Brookhaven Instruments).

### Airway epithelial cell mucus collection

Airway epithelial cell cultures were washed with phosphate buffered saline (PBS) to remove the accumulated mucus from the apical surface for collection. The collected mucus was filtered using Amicon ultra centrifugal filter units with a 100 kDa cutoff to remove excess PBS. The resulting mucus was stored at 4 °C until time of use. Mucus collected from NHBE cells was pooled for all experiments.

### Disulfide bond concentration assay

For mucus collected from Calu-3, BCi-NS1.1, and NHBE cells, the disulfide bond concentration was determined using a previously established protocol.^40^ Briefly, samples were resuspended in 8 M Guanidine-HCl to bring the final volume to 500 μL before treatment with 10% (v/v) 500 mM iodoacetamide at room temperature for 1 hour. Samples were subsequently treated with 10% (v/v) 1 M DTT at 37 °C for 2 hours. Small molecules were removed by passing samples through 7 k MWCO Zebra desalting columns, which were also used to exchange the buffer for 50 mM Tris-HCl (pH 8.0). Equal volumes of sample and 2 mM monobromobimane were combined in a 96-well plate before incubation at room temperature for 15 minutes. The samples were read at 395 nm excitation and 490 nm emission and compared to a standard curve of L-cysteine to determine the disulfide bond concentration.

### Mucin content assay

Using a previously established protocol,^41^ mucus collected from Calu-3, BCi-NS1.1, and NHBE cells was analyzed to determine the relative mucin content. Briefly, 50 μL of samples were combined with 60 μL of alkaline CNA reagent, which was made by combining 200μL of 0.6 M 2-cyanoacetamide and 1 mL of 0.15 M NaOH. Samples were then incubated at 100 °C for 30 minutes. Subsequently, 0.5 mL of 0.6 M borate buffer, pH 8.0, was added to each sample. After samples cooled for 15 minutes at room temperature, the fluorescence intensity was measured at 336 nm excitation and 383 nm emission. Mucin from porcine gastric mucin was dissolved in mucin buffer and analyzed with this protocol to generate a standard curve. Mucin buffer was made by combining 0.01 M Na_2_HPO_4_ and 0.04% NaN_3_ at pH 7.4.

### Sialic acid concentration assay

Sialic acid (SA) concentration was determined in mucus samples using a SA assay kit (Sigma-Aldrich, MAK314) and following the manufacturer protocol. Briefly, samples were hydrolyzed to release bound SA. These hydrolyzed samples were used to determine the total SA concentration. Samples were then combined with thiobarbituric acid. Samples were oxidized, resulting in the oxidation of SA into formylpyruvic acid, which reacts with thiobarbituric acid to form a pink product. The fluorescence of the product for each sample was measured at 555 nm excitation and 585 nm emission and compared to a standard curve of SA to determine the SA concentration in each sample.

### Fluorescence imaging and multiple particle tracking analysis

To evaluate the movement of particles in HAE mucus, 1 μL of each type of particle was added to 20 μL of HAE mucus and placed on a slide in the middle of a vacuum grease-coated O-ring. Slides were equilibrated for 30 minutes at room temperature prior to fluorescence imaging with a Zeiss Confocal LSM 800 microscope equipped with a 63x water-immersion objective. Multiple 10-second videos were recorded at 33.3 frames per second for each sample. Fluorescence microscopy video files were processed using a previously developed MATLAB code capable of tracking multiple particles and calculating the MSD. The MSD was calculated as ⟨*MSD*(*τ*)⟩ = ⟨(*x*^2^ + *y*^2^)⟩, for each particle.^42–44^ The MSD values for NP were then used to calculate the microrheological properties using the generalized Stokes-Einstein relation,^45^ as 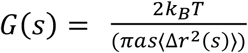 gives the viscoelastic spectrum where *k*_*B*_*T* is the thermal energy, *a* is the radius, and *s* is the complex Laplace frequency.^39^ The complex modulus *G*\* was calculated as *G*^*\**^(*ω*) = *G*^*’*^(*ω*) + *G*^*”*^(*iω*) where *iω* is used in place of *s, i* is a complex number, and *ω* is the frequency^18^. The pore size (*ξ*) can be estimated from the *G’* values as 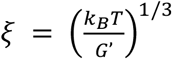.^39^ Using the previously published machine learning-based analysis and the particle survival analysis,^24,25,27^ the trajectory data for each viral strain and the muco-inert nanoparticles were analyzed to classify particle movement as fractional Brownian motion (FBM), Brownian motion. (BM), or continuous time random walk (CTRW). The resulting classified trajectories were used in the particle survival analysis to determine the percentage of particles that were predicted to cross the mucosal barrier.

### Enzymatic and chemical alteration of mucus glycans

Collected NHBE mucus was incubated with 10 μL exogenous α2-3,6,8,9-Neuraminidase (NA_ex_) from *Arthrobacter ureafaciens* (5 U/mL, Millipore Sigma) in 5 mL of reaction buffer (0.1 M sodium acetate, pH 5.5) for 2 hours at 37 °C. The treated samples were washed with PBS to remove NA_ex_ and biproducts before subsequent concentration via Amicon ultra centrifugal filter units with 100 kDa MW cutoff. Samples were then resuspended to original volume with PBS. Mucus collected from NHBE cells was incubated in 5 mL of reaction buffer (0.1 M sodium acetate, pH 5.5) with 2 mM ice-cold sodium periodate (NaIO_4_) for 30 minutes at 4 °C while protected from light. Unreacted sodium periodate was quenched with excess ethylene glycol. The treated samples were washed with PBS to remove reactant materials before subsequent concentration via Amicon ultra centrifugal filter units with 100 kDa MW cutoff. Concentrated samples were resuspended to original volume with PBS.

### IAV challenge in BCi mucus and PEG gel coated HAE Cultures

Fully differentiated BCi-NS1.1 cells were cultured as described above, washed to remove native mucus, and divided into 3 treatment groups (3 cultures per condition): untreated (i.e. no mucus added post-wash), cells transplanted with BCi mucus post-washing, and cells transplanted with a PEG-hydrogel post-washing. BCi mucus transplanted onto the washed cultures was collected from mature HAE cultures at ALI, concentrated using a 100k amicron filter, and stored at -80°C until usage. Polyethylene glycol (PEG) hydrogels solution components, 4% (w/v) polyethylene glycol orthopyridyl disulfide (PEG-OPSS, 5kD, Creative PEGWorks) and 4% (w/v) polyethylene glycol thiol (PEG-SH, 10kD, LaysenBio) were individually prepared in PBS and then sterilized with a 0.45 μm syringe filter. Cultures were then treated apically with either 20uL of BCi mucus or 10uL of 4% w/v PEG-OPSS solution followed by 10uL of 4% w/v PEG-SH solution which rapidly form into a PEG hydrogel on the transwell. Mucus and PEG hydrogel were dispersed through gentle shaking followed by a 50-minute equilibration period at 37°C. A small 4 μL volume of Udorn IAV (1.5 x 10^4^ pfu per well for MOI of ∼0.1) was then centrally administered to HAE cultures and incubated for 15 minutes at 37°C. Following inoculation, cultures were washed twice with PBS for 10 minutes at 37°C to remove the hydrogel coatings and non-absorbed virus. To prevent secondary infection, basolateral media was replaced with fresh media containing zanamivir (1.25 μM) and the apical compartment was washed with PBS containing zanamivir (1.25 μM). Twelve hours post infection, cultures were fixed in 4% paraformaldehyde solution before permeabilization in 2.5% Triton X-100. Cells were then blocked in 3% BSA before adding primary antibody mouse monoclonal anti-IAV nucleoprotein A1 and A3 blend antibody (EMD Millipore, MAB8251). The primary antibody was followed by a secondary anti-mouse Alexa-488 (Invitrogen) antibody, and then DAPI (Invitrogen) for nuclei staining. A series of 5 images per well taken at 10×magnification using a Zeiss LSM 800 Confocal microscope and the percentage of the culture area successfully infected was quantified using ImageJ.

### Statistical Analysis

Data were statistically analyzed using GraphPad Prism 10 (GraphPad Software, San Diego, CA). Specific analyses used for each data set are noted in the figure captions.

## Notes

### Competing Interest Statement

The authors have declared no competing interest.

### Summary of Updates

All prior figures and manuscript text have been revised to include newly acquired data.

