## Supplemental Information for "Mucus physically restricts influenza A viral particle access to the epithelium"

| Donor | Age | Sex | Race |
|---|---|---|---|
| 1 | 65 | Male | Caucasian |
| 2 | 73 | Female | Caucasian |
| 3 | 52 | Male | Hispanic or Latino |
| 4 | 65 | Female | Caucasian |

**Supplemental Table 1.** Donor information for NHBE cells.

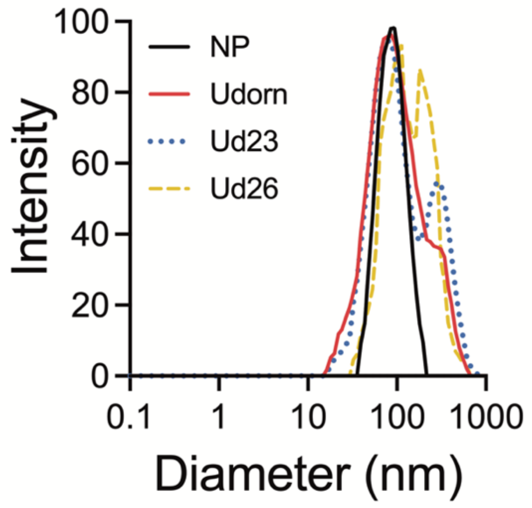


**Figure S1. Particle size of NP and IAV.** Measured hydrodynamic radius of NP, Udorn, Ud23, and Ud26. NP is shown as black solid line, Udorn is shown as red solid line, Ud23 is shown as blue dotted line, and Ud26 is shown as yellow dashed line.

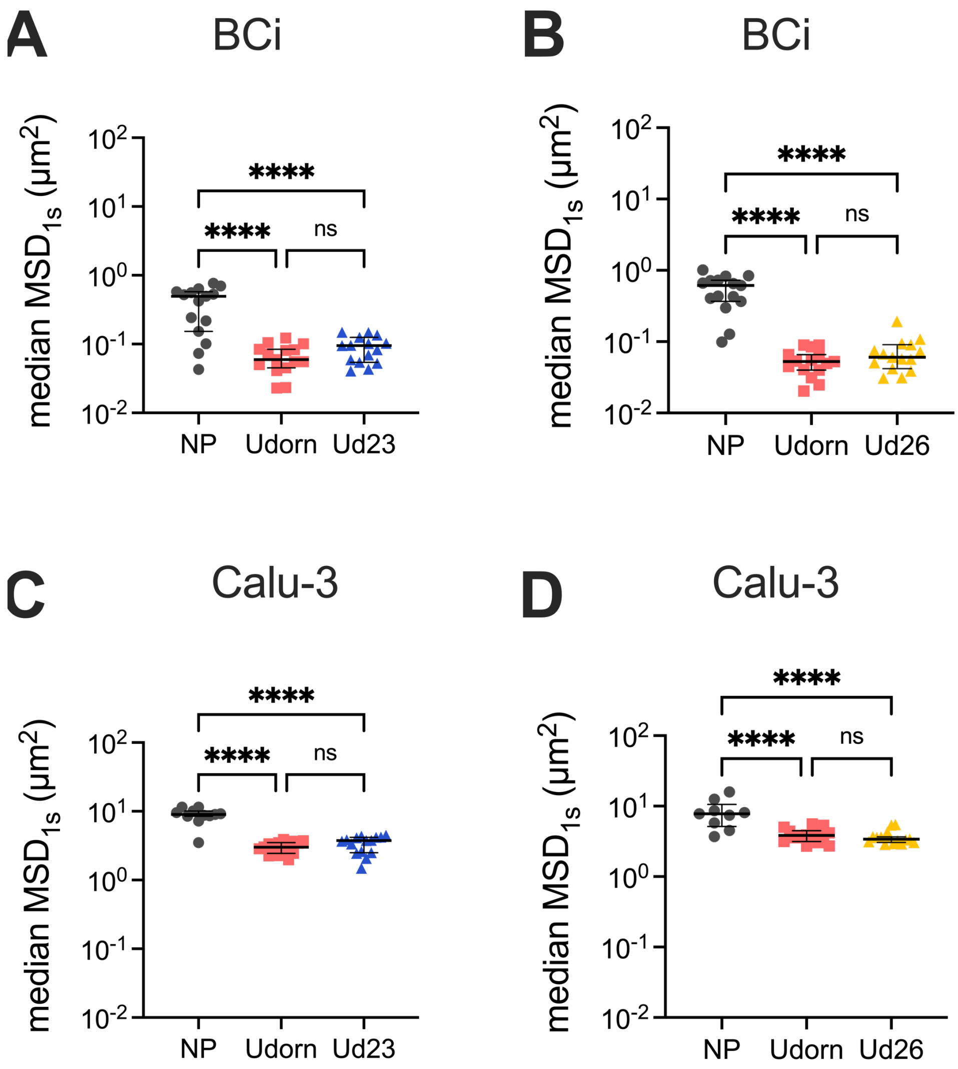


**Figure S2. Diffusion of wildtype and mutant Udorn IAV within Calu-3 and BCi mucus. (A)** Median $MSD_{1s}$ for NP, Udorn, and Ud23 diffusing in BCi mucus. (**B**) Median $MSD_{1s}$ for NP, Udorn, and Ud26 diffusing in BCi mucus. **(A)** Median $MSD_{1s}$ for NP, Udorn, and Ud23 diffusing in Calu-3 mucus. (**B**) Median $MSD_{1s}$ for NP, Udorn, and Ud26 diffusing in Calu-3 mucus. NP is shown as black circles, Udorn is shown as red squares, Ud23 is shown as blue triangles, and Ud26 is shown as yellow triangles.

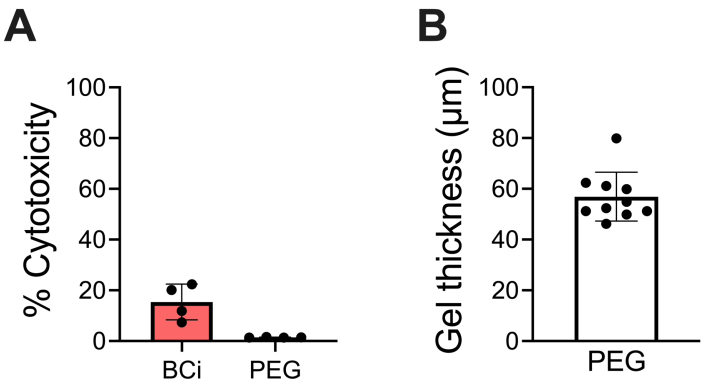


**Figure S3. PEG hydrogel biocompatibility and thickness following apical application on HAE cultures.** (**A**) Cytotoxicity was evaluated in HAE cultures treated apically with either 30 µL of BCi mucus or 30 µL of PEG hydrogels via LDH assay. Following 2 hours of incubation, BCi mucus and PEG gels were removed through a PBS wash. Washed uncoated HAE cultures were treated with lysis buffer diluted 1:10 in PBS for 45 minutes at 37ºC to determine maximum (100%) LDH activity. PBS was apically added to all cultures (n=4 per condition) and incubated at 37°C for 30 minutes. LDH activity in apical washings was determined using Promega CytoTox 96TM Nonradioactive Cytotoxicity Assay (Promega G1780). **(B)** To measure the height of the PEG hydrogel following apical application on transwells, 4% PEG-SH (10 kDa) and 4% PEG-OPSS (5 kDa) with Texas Red dextran (10kD, 2 mg/mL) solutions were individually added to the transwell to allow for the formation of the hydrogel on the transwells. The hydrogel was dispersed over the membrane through gentle shaking. Following gel solidification, transwell membranes were cut from the casing and mounted on a coverslip. Images of the height were obtained through five Z-stack images, one centrally and four circumferential, using a 10X objective lens. Hydrogel height was measured using a custom ImageJ macro capturing 90% gray area intensity.

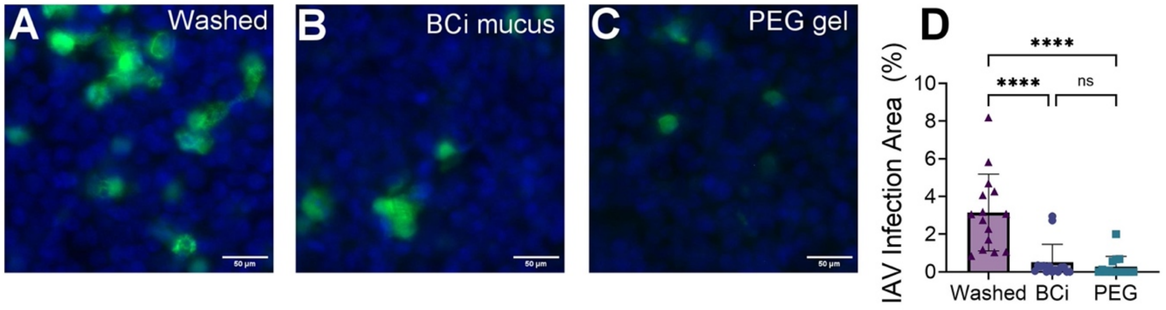


**Figure S4. IAV infection in human airway epithelial cultures with mucus and synthetic PEG gel coatings. (A-C)** Fluorescence micrographs of (**A**) washed, (**B**) BCi mucus coated, and (**C**) PEG gel coated HAE cultures infected apically with Udorn IAV 12 hpi after 15 min of inoculation. Green indicates staining for IAV nucleoprotein and blue indicates DAPI-stained nuclei. Scale bar = 50 µm. (**D**) Percentage of HAE culture area infected as determined by IAV nucleoprotein staining. Each data point representative of individual fluorescent micrographs collected from 3 individual cultures per experimental condition. For these experiments, HAE cultures were not treated with zanamavir to prevent secondary infection as in the data shown Figure 5E-H.

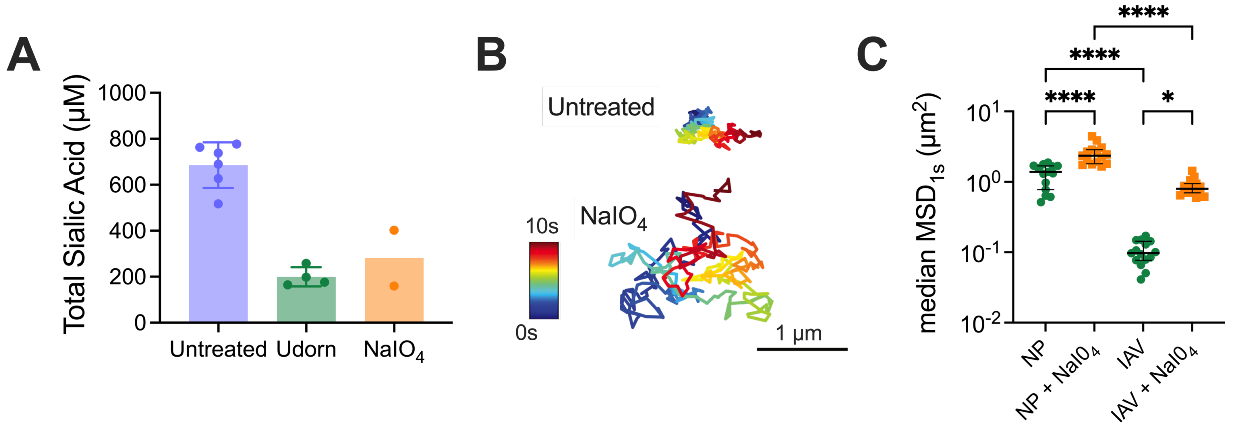


**Figure S5. Chemical alteration of mucus glycans and IAV diffusion. (A)** Representative trajectories of Udorn in untreated and $NaIO_4$ treated NHBE mucus. Trajectory color changes with time, with dark blue indicating 0 s and dark red indicating 10 s. Scale bar = 1 μm. **(B)** Characterization of total Sia concentration for untreated, Udorn treated, and $NaIO_4$ treated NHBE mucus. Lines drawn to indicate levels of Sia measured in Calu-3 and BCi samples from Figure 1. **(C)** Median $MSD_{1s}$ for NP and Udorn in untreated and $NaIO_4$ treated NHBE mucus. Data set in **(C)** analyzed with one-way analysis of variance (ANOVA) and Šídák's multiple comparisons test: $^*p < 0.05$, $^{****}p < 0.0001$.
